# Olfactory rod cells: a rare cell type in the larval zebrafish olfactory epithelium with an actin-rich apical projection

**DOI:** 10.1101/2020.11.04.367979

**Authors:** King Yee Cheung, Suresh J. Jesuthasan, Sarah Baxendale, Nicholas J. van Hateren, Mar Marzo, Christopher J. Hill, Tanya T. Whitfield

**Author notes:** Authors for correspondence: Tanya T. Whitfield, Suresh J. Jesuthasan.

## Abstract

We report the presence of a rare cell type, the olfactory rod cell, in the developing zebrafish olfactory epithelium. These cells each bear a single actin-rich rod-like apical projection extending about 10 μm from the epithelial surface. Live imaging with a ubiquitous Lifeact-RFP label indicates that the rods can oscillate. Olfactory rods arise within a few hours of the olfactory pit opening, increase in numbers and size during larval stages, and can develop in the absence of olfactory cilia. Olfactory rod cells differ in morphology from the known classes of olfactory sensory neuron, but express reporters driven by neuronal promoters. The cells also differ from secondary sensory cells such as hair cells of the inner ear or lateral line, or sensory cells in the taste bud, as they are not associated with established synaptic terminals. A sub-population of olfactory rod cells expresses a Lifeact-mRFPruby transgene driven by the *sox10* promoter. Mosaic expression of this transgene reveals that olfactory rod cells have rounded cell bodies located apically in the olfactory epithelium.

## Introduction

The vertebrate olfactory epithelium (OE) enables the detection of chemical cues, giving rise to the sense of smell (reviewed in [Axel, 1995]). The function of this epithelium, which derives from paired cranial neurogenic placodes, is mediated by a diverse set of cells that includes neuronal receptors and non-sensory cells. Olfactory sensory neurons (OSNs) are bipolar neurons that extend a dendrite to the apical surface of the OE, and an axon to the olfactory bulb (OB). In mammals, two broad classes of sensory receptors — ciliated and microvillous OSNs — have been identified on the basis of morphology, receptor expression and OB target. Mammalian OSNs can act as both chemosensors and mechanosensors (Grosmaitre et al., 2007; Iwata et al., 2017). The OE of other vertebrates also contains ciliated and microvillous neurons. In fish, additional classes of OSNs have been identified. Each occupies a stereotyped position within the pseudostratified OE, with the dendrite bearing a distinct and characteristic specialisation projecting into the environment (Hansen & Zeiske, 1998; Hansen & Zielinski, 2005; Sato et al., 2005; reviewed in [Maier et al., 2014]).

In zebrafish, ciliated neurons, which express olfactory marker protein (OMP) and odorant receptor (OR) genes, have a cell body that lies deep within the OE, an axon that projects to dorsal and medial regions of the OB, and a slender dendrite extending to the surface of the olfactory pit. Here, the dendritic knob bears a cluster of primary cilia that project into the olfactory cavity (Hansen & Zeiske, 1998; Hansen & Zielinski, 2005; Sato et al., 2005). Microvillous neurons, characterised by the expression of TrpC2 and vomeronasal (VR)-type pheromone receptors, have cell bodies that lie in the intermediary layer of the OE, an axon that projects to the lateral part of the OB, and a dendrite bearing a tuft of short, actin-rich microvilli (Hansen & Zeiske, 1998; Hansen & Zielinski, 2005; Sato et al., 2005). Crypt neurons, less abundant than ciliated or microvillous neurons, have rounded cell bodies that sit apically in the OE, with both cilia and microvilli extending from a crypt within the cell body (Hansen & Zeiske, 1998; Hansen & Zielinski, 2005; Parisi et al., 2014; Biechl et al., 2016; Bettini et al., 2017). Kappe neurons lie in the superficial layers of the adult zebrafish OE and are named for their apical actin-rich cap, presumed to be microvilli (Ahuja et al., 2014). Pear-shaped neurons are also positioned superficially in the adult OE and have short apical dendrites, but express some markers in common with ciliated neurons (Wakisaka et al., 2017).

OSNs are surrounded and separated by a network of non-neuronal cell types. These include sustentacular (support) cells, basal cells that replenish the OSNs, and goblet cells that produce mucus (Hansen & Zeiske, 1993; Hansen & Zeiske, 1998; reviewed in [Olivares & Schmachtenberg, 2019]; Demirler et al., 2019). In fish, multiciliated cells, located around the rim of the olfactory pit, each bear multiple long motile cilia. These have a characteristic 9+2 axoneme and beat at around 24 Hz, resulting in an asymmetric flow that draws water and odorants into the olfactory cavity and flushes them out again (Reiten et al., 2017).

In addition to common cell types, tissues may also contain rare or sparsely-distributed cell types, which are difficult to detect by conventional histological methods. These include stem cells, immune cells, or other rare cell types, which can have critical functions (see, for example, [Montoro et al., 2018; Sui et al., 2018]). Characterisation of the identity and lineage of every cell type of an organ system is a goal of many contemporary single-cell and single-nucleus RNA-seq studies (Junker et al., 2014; Satija et al., 2015; Hernández et al., 2018; Raj et al., 2018; Farnsworth et al., 2020; Wattrus & Zon, 2020). Transgenic or other fluorescent markers, coupled with high-resolution imaging of the whole embryo, can also help to identify cell types that may previously have been overlooked (see, for example, [Kawakami et al., 2010; Galanternik et al., 2017; van Lessen et al., 2017]).

We report here the existence of a rare cell type, the olfactory rod cell, in the developing zebrafish OE. Olfactory rod cells are characterised by a single actin-rich apical projection. The morphology of the rod matches brief descriptions of similar structures in the OE of several other fish species, many of which were previously dismissed either as senescent forms of OSNs or as fixation artefacts. Using a variety of imaging techniques and transgenic lines, including live imaging, we show that zebrafish olfactory rod cells are present in living fish and can be detected from early stages of larval development.

## Materials and Methods

### Zebrafish husbandry

Zebrafish strains used in this study were wild type (AB strain – ZFIN), *ift88*^*tz288b*^ (Tsujikawa & Malicki, 2004), *sox10*^*m618*^ (Dutton et al., 2001), *Tg(actb2:Lifeact-RFP)*^*e115*^ (Behrndt et al., 2012), *Tg*(*actb2:Lifeact-GFP*)^*e114*^ (Behrndt et al., 2012), *Tg(Xla.Tubb:jGCaMP7f)*^*sq214*^ (Chia et al., 2019), *Tg(elavl3:GCaMP6f)*^*jf1*^ (Dunn et al., 2016), *Tg(elavl3:H2B-GCaMP6s)*^*jf5*^ (Dunn et al., 2016)*, Tg(pou4f3:GAP-GFP)*^*s356t*^ (Xiao et al., 2005) and *Tg(sox10:Lifeact-mRFPruby)*^*sh630*^ (this study). Homozygous *sox10*^−/−^ mutant larvae were identified by their lack of body pigmentation at 5 days post-fertilisation (dpf). Adult zebrafish were kept in a 10 hours dark/14 hours light cycle at 28.5°C and spawned by pair-mating or marbling (Aleström et al., 2019). Eggs were collected and staged according to standard protocols (Kimmel et al., 1995; Nüsslein-Volhard & Dahm, 2002), and raised in E3 medium (5 mM NaCl, 0.17 mM KCl, 0.33 mM CaCl_2_, 0.33 mM MgSO_4_, with 0.0001% methylene blue at early stages) at 28.5°C. For controlling the developmental rate to obtain embryos at stages 34-46 hours post-fertilisation (hpf), embryos were incubated at 25°C or 34°C in accordance with Kimmel’s formula, *H*_*T*_ = *h* ÷ (0.055*T* − 0.57) (Kimmel et al., 1995). For live imaging, zebrafish were anaesthetised with 0.5 mM tricaine mesylate in E3.

### Generation of the Tg(sox10:Lifeact-mRFPruby) transgenic line

The *-4725sox10:Lifeact-mRFPruby* construct was generated using the Gateway Tol2 kit (Kawakami, 2007; Kwan et al., 2007). The p5E *-4725sox10* promoter (Dutton et al., 2008; Rodrigues et al., 2012), pME-*Lifeact-mRFPruby* (Riedl et al., 2008), and p3E polyA sequences were cloned into pDestTol2pA3 through an LR Clonase reaction. The 12.1 kb final plasmid was sequenced and injected into the AB strain. Injected embryos were grown to adulthood and crossed to AB. Transgenic progeny from one founder male were selected based on mRFPruby expression in the inner ear and grown to adulthood to generate a stable line. Embryos with bright fluorescence, presumed to be homozygous for the transgene, were chosen for imaging.

### Immunohistochemistry and phalloidin staining

Zebrafish embryos and larvae were fixed in 4% paraformaldehyde (PFA) in phosphate-buffered saline (PBS) for two hours at room temperature or overnight at 4°C. Zebrafish were washed three or more times with PBS, and permeabilised by incubation in PBS-Triton X-100 (0.2% Triton for 32-48 hpf embryos, 1% Triton for later stages) for several hours at 4°C until staining.

To visualise F-actin, zebrafish were stained with either Alexa Fluor 488 phalloidin (Cell Signaling Technology; 1:150), Alexa Fluor 568 (Invitrogen ThermoFisher; 1:50), or Alexa Fluor 647 phalloidin (Invitrogen ThermoFisher; 1:50) in PBS overnight at 4°C. After staining, zebrafish were washed four times in PBS over two or more hours before imaging.

For antibody staining, after fixing and washing, zebrafish were washed a further three times in PBS-0.2% Triton and incubated in blocking solution (10% sheep serum in PBS-0.2% Triton for acetylated α-tubulin staining; 1% bovine serum albumin (BSA) in PBS-0.1% Triton for SV2 staining) for 60 minutes at room temperature. Primary antibodies were mouse IgG1 anti-acetylated α-tubulin antibody (Sigma-Aldrich; 1:100) and mouse IgG1 anti-SV2 antibody (deposited in Developmental Studies Hybridoma Bank by K. M. Buckley; 1:100). Staining was carried out in blocking solution containing 1% dimethyl sulfoxide (DMSO; Sigma-Aldrich) overnight at 4°C. Zebrafish were washed three times in PBS-0.2% Triton, and a further four times over two or more hours. The secondary antibody was Alexa 647-conjugated goat anti-mouse IgG1 (Invitrogen ThermoFisher; 1:200). For double stains with phalloidin, Alexa Fluor 488 phalloidin (1:150) and DMSO (1%) were added together with the secondary antibody in blocking solution overnight at 4°C. Zebrafish were then washed four times in PBS-0.2% Triton and stored at 4°C until imaging. Controls with no primary antibody yielded no staining (not shown).

### Ototoxin treatment

For neomycin treatment, a concentration of 500 μM was chosen, as it was an effective concentration used by Harris et al. (2003) for minimum lateral line hair cell survival, as measured by DASPEI staining. A 5 mM solution was made by adding neomycin trisulfate salt hydrate (Sigma-Aldrich) to MilliQ water and used at a 1:10 dilution in E3 fish medium. *Tg(pou4f3:GFP)* transgenic zebrafish were treated for 60 minutes at 28.5°C. An equivalent volume of MilliQ water in E3 was used for the control group. Zebrafish were washed three times in fresh E3 and left at 28.5°C for two hours. GFP signal was screened using widefield fluorescence microscopy to analyse hair cell damage. Zebrafish were fixed and stained with Alexa Fluor 647 phalloidin as above.

### Fluorescence imaging

For confocal imaging, fixed zebrafish embryos and larvae were mounted in 1.5% low melting point (LMP) agarose in PBS, and live zebrafish were mounted in 1.5% LMP agarose in E3 in WillCo glass-bottomed dishes (mounted in frontal view for 32-48 hpf, dorsal view for later stages). Zebrafish were imaged on a Zeiss LSM880 Airyscan confocal microscope equipped with a Plan-Apochromat 20×/0.8 M27 air objective, LD LCI Plan-Apochromat 40×/1.2 Imm Korr DIC M27 water immersion objective, or Plan-Apochromat 63×/1.4 oil DIC M27 objective. Images were acquired in Airyscan SR mode, Airyscan Fast scan mode with SR sampling, or Airyscan Fast scan mode with Opt sampling. Zebrafish were also imaged on a Zeiss LSM 800 attached to an upright microscope with a W Plan-Apochromat 40×/1.0 DIC M27 or 63×/1.0 M27 water dipping objective. The laser lines used were 488, 561, and 633 nm. Widefield imaging was performed on a Zeiss Axio Zoom.V16 fluorescence stereo zoom microscope equipped with a Zeiss 60N-C 1” 1.0× C-mount and AxioCam MRm camera. For fast-capture time series imaging, live zebrafish larvae were mounted in 0.9% LMP agarose in E3 and imaged on a Zeiss Z1 Light-sheet microscope, with 4% tricaine in E3 in the sample chamber. Imaging was performed with a W Plan-Apochromat 20× objective using brightfield illumination and the 561 nm laser line. Images were acquired at a rate of 50.07 frames per second (fps).

### Scanning electron microscopy

For scanning electron microscopy, *ift88* homozygous mutant and phenotypically wild-type sibling larvae at 4 dpf were fixed overnight in 2.5% glutaraldehyde/0.1M sodium cacodylate buffer. Samples were washed in buffer, post-fixed in 2% aqueous osmium tetroxide for 1 hour, washed in buffer again and then dehydrated through a graded ethanol series (50%, 75%, 95%, 100%) before being dried in a mixture of 50% hexamethyldisilazane (HMDS) in 100% ethanol. Final drying was in 100% HMDS. After removal of the final HMDS wash, samples were left to dry in a fume hood overnight. Samples were mounted onto a pin stub using a Leit-C sticky tab and Leit-C mounting putty, gold-coated using an Edwards S150B sputter coater, and examined in a Tescan Vega3 LMU Scanning Electron Microscope at an operating voltage of 15 kV and imaged using a secondary electron detector.

### Image processing, quantification, and statistical analyses

Zeiss LSM880 Airyscan confocal images were subjected to Airyscan processing on Zen Black 2.3 software (Zeiss) using “Auto” Airyscan processing parameters. Further processing was performed on Fiji (Schindelin et al., 2012). 3D rendering was performed using the 3D Viewer plugin (Schmid et al., 2010) on Fiji. Rod projection lengths were measured in 3D from confocal images using Fiji, and calculated in Microsoft Excel using the PyT method (based on the Pythagorean theorem) from Dummer et al. (2016). All quantifications were exported into GraphPad Prism 8, which was then used for performing statistical analyses and making graphs.

Statistical analyses were carried out in GraphPad Prism 8. Datasets were considered normally distributed if they passed at least one of four normality tests (Anderson-Darling, D’Agostino & Pearson, Shapiro-Wilk, and Kolmogorov-Smirnov tests). Statistical tests used are stated in the figure legends. Bars on graphs indicate mean ± standard error of the mean (S.E.M.), unless stated otherwise. *P* values are indicated as follows: *P* > 0.05 (not significant, ns), *P* < 0.05 (*), *P* < 0.01 (**), *P* < 0.001 (***), *P* < 0.0001 (****).

For mapping spatial distributions of rod cells within the olfactory pit, 2D maximum intensity projection images were imported into the Desmos Graphing Calculator (desmos.com). The positions and sizes of the images were adjusted to align the rims of olfactory pits with an ellipse to fit the shape of the rim, defined by 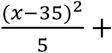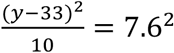. The positions of the base of each rod, relative to the ellipse, were plotted as coordinates onto the graph. The resulting graphs were exported as .png image files.

Figures were prepared using Adobe Photoshop.

## Results

### Actin-rich rod-like apical projections, distinct from microvilli and cilia, are present in the olfactory epithelium of larval and juvenile zebrafish

Staining of the wild-type larval and juvenile zebrafish OE with fluorescently-conjugated phalloidin, which binds to F-actin, reveals the presence of several actin-rich rod-like projections (‘rods’) in each olfactory pit (Figure 1A-B’). These projections differ in number, distribution, size and morphology from any of the described apical projections of zebrafish OSNs. The projections extend from below the apical surface of the OE and project about 10 μm above it, tapering to a point. This is an order of magnitude longer than OSN microvilli, which are typically 0.5-0.8 μm in length (Hansen & Zeiske, 1998). Olfactory rods are shorter than the surrounding phalloidin-negative olfactory cilia (Fig. 1C-D’), and do not label with an anti-acetylated α-tubulin antibody (Figure 1C-C’’’). Rods are not evenly distributed across the OE, but are mostly clustered posterolaterally in each olfactory pit, although there is variation between individuals (Figure 1E). At low magnification, the olfactory rods appear similar to the actin-rich stereociliary bundle of mechanosensory hair cells of the inner ear and lateral line. However, higher magnification images reveal that the olfactory rod is not oligovillous, but appears to be a single structure (Figure 1B’, C’’’, D’). This contrasts with the stepped array of multiple stereocilia present on the apical surface of mechanosensory hair cells (Figure 1F).

**Figure 1.**
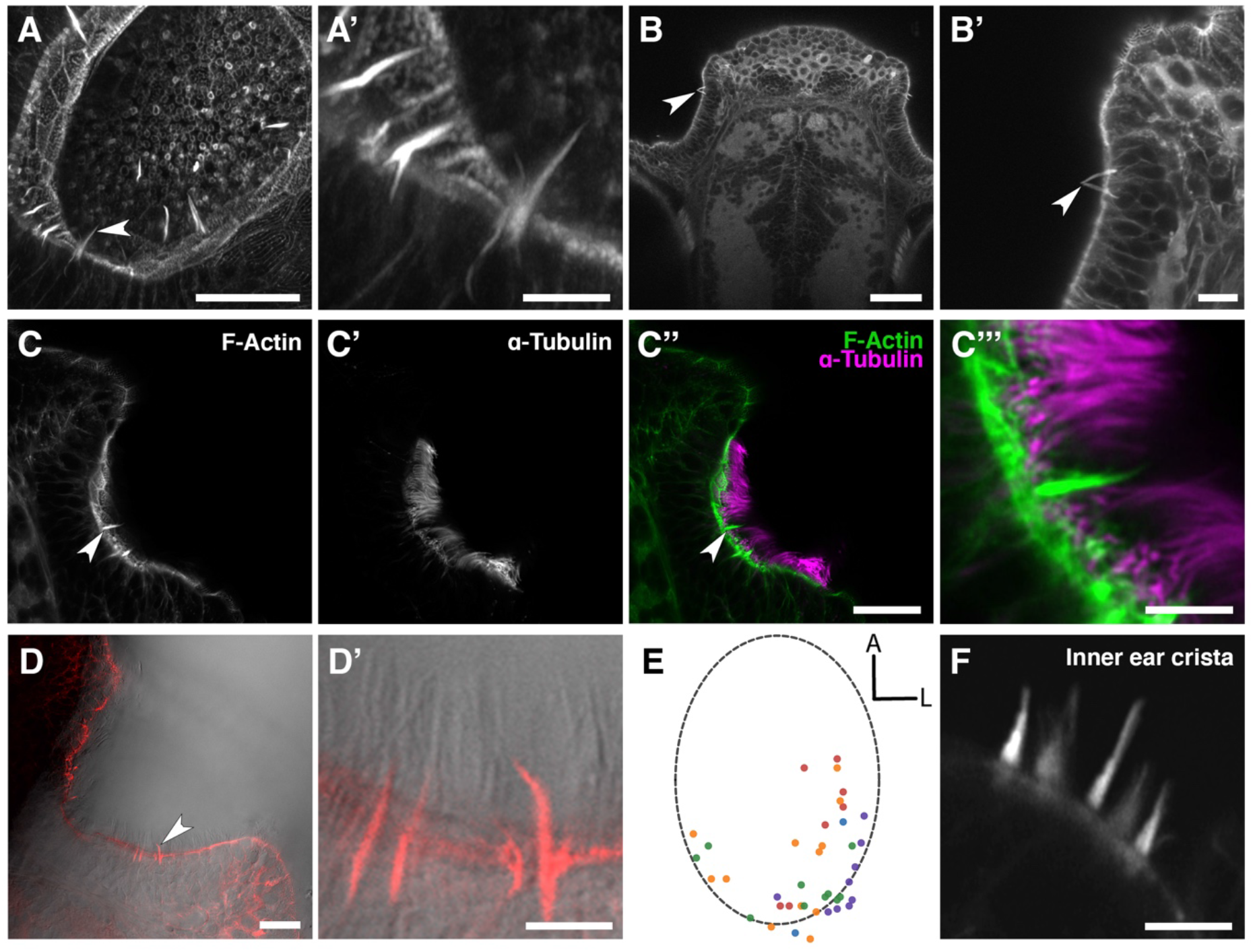
Phalloidin staining reveals the presence of actin-rich rod-like projections, distinct from microvilli and cilia, in the zebrafish larval and juvenile olfactory epithelium. (A) Maximum intensity projection of an Airyscan confocal image of phalloidin stain in an olfactory pit of a 5 dpf wild-type larva; anterior to the top right, lateral to the bottom right. Arrowhead marks one example olfactory rod. Scale bar = 20 μm. (A’) Enlargement of olfactory rods in A. Scale bar = 5 μm. (B) Dorsal view low power image of phalloidin stain in the head of an 18 dpf (5 mm) wild-type juvenile zebrafish; anterior to the top. Arrowhead marks the position of two olfactory rods in an olfactory pit. Scale bar = 50 μm. (B’) Enlargement of OE in B. Arrowhead marks two olfactory rods. Scale bar = 10 μm. (C-C’’) Airyscan confocal image of Alexa-phalloidin signal (C), acetylated α-tubulin immunohistochemistry signal (C’), and merged signals (C’’) in an olfactory pit of a 4 dpf wild-type larva; anterior to the top, lateral to the right. Arrowhead marks one example olfactory rod. Scale bar = 20 μm. (C’’’) Enlargement of olfactory rod in C’’. Scale bar = 5 μm. (D) Differential interference contrast (DIC) image and phalloidin stain (red) in an olfactory pit of a 5 dpf wild-type larva; anterior to the top, lateral to the right. Arrowhead marks one example olfactory rod. Scale bar = 20 μm. (D’) Enlargement of olfactory rods in D. Surrounding olfactory cilia are visible and unlabelled by Alexa-phalloidin. Scale bar = 5 μm. (E) A map of the positions of olfactory rod cell projection bases in olfactory pits of 4 dpf wild-type larvae (*N* of olfactory pits = 5), based on 2D maximum intensity projections of confocal images of phalloidin stains; anterior ‘A’ to the top, lateral ‘L’ to the right. One dot represents one olfactory rod. Different coloured dots represent rods from different larvae. (F) Airyscan confocal image of phalloidin stain in an inner ear crista of a 5 dpf wild-type larva. Hair cell stereocilia are labelled with Alexa-phalloidin, and are arranged in a stepped array. In the stereociliary bundle on the extreme left, four different stereociliary lengths are visible (compare with A’). Scale bar = 5 μm.

To characterise the timing of appearance and development of the olfactory rods during embryonic and larval stages, we stained fixed samples from 36 hpf, just after formation of the olfactory pits (Hansen & Zeiske, 1993), to 5 dpf. Occasional rods were present in olfactory pits at 36 hpf, but were only consistently present beyond 46 hpf (Figure 2A, B). Although the number of rods per olfactory pit varied at each stage, the average number increased over time. By 5 dpf, each olfactory pit contained 10.7 ± 2.9 (mean ± standard deviation, s.d.) rods (Figure 2B). After measuring the rods in 3D, we found an increase in projection length (from the base of the phalloidin-positive projection to the tip) from 36 hpf to 5 dpf, with the most significant increase occurring by 48 hpf, despite a relatively large range in length at each stage. At 5 dpf in fixed samples, the mean projection length was 10.4 ± 2.2 (s.d.) μm, with the largest measuring 17.5 μm (Figure 2C).

**Figure 2.**
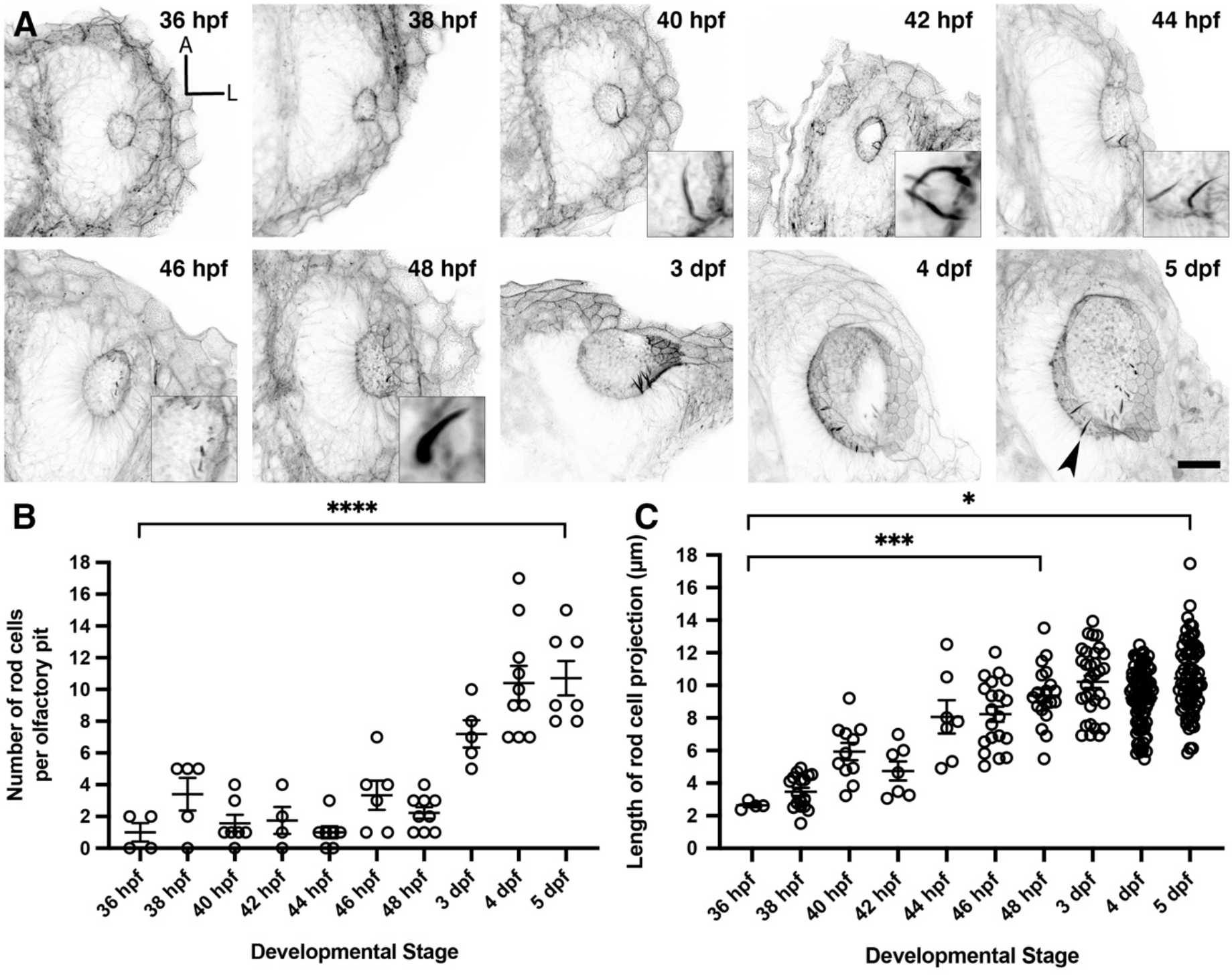
Olfactory rod cells arise early during zebrafish olfactory pit development. (A) Maximum intensity projections of Airyscan confocal images showing the wild-type development of olfactory pit and olfactory rod cells at various embryonic and larval stages, using Alexa-phalloidin as a marker; anterior ‘A’ to the top, lateral ‘L’ to the right. Grayscale values from the original fluorescence image have been inverted. Arrowhead marks one example olfactory rod. Scale bar = 20 μm. Selected inserts show olfactory rods at higher magnification. (B) The change in number of olfactory rod cells per olfactory pit during embryonic development – 36 hpf (*N* of olfactory pits = 4), 38 hpf (*N* = 5), 40 hpf (*N* = 7), 42 hpf (*N* = 4), 44 hpf (*N* = 7), 46 hpf (*N* = 6), 48 hpf (*N* = 9), 3 dpf (*N* = 5), 4 dpf (*N* = 10), and 5 dpf (*N* = 7). Bars indicate mean ± S.E.M. for each stage. Linear regression analysis; **** indicates *P* < 0.0001. (C) The change in lengths of olfactory rod cell projections during embryonic development – 36 hpf (*N* of olfactory pits = 2, *n* of olfactory rods = 4), 38 hpf (*N* = 4, *n* = 17), 40 hpf (*N* = 6, *n* = 11), 42 hpf (*N* = 3, *n* = 7), 44 hpf (*N* = 5, *n* = 7), 46 hpf (*N* = 6, *n* = 20), 48 hpf (*N* = 9, *n* = 20), 3 dpf (*N* = 5, *n* = 32), 4 dpf (*N* = 10, *n* = 82), and 5 dpf (*N* = 8, 71). Bars indicate mean ± S.E.M. for each stage. Linear regression analysis; * indicates *P* = 0.0251, *** indicates *P* = 0.0009.

### Olfactory rod cell projections can develop in the absence of olfactory cilia

As described above, olfactory rods differ from olfactory cilia in terms of size, shape, cytoskeletal composition, and distribution in the OE. We therefore hypothesised that olfactory rod cell projections would not be affected by mutations that disrupt the formation of cilia. To test this, we examined fish mutant for *ift88*, which codes for a component of the intraflagellar transport machinery necessary for the normal formation and maintenance of cilia (Tsujikawa & Malicki, 2004). A phalloidin stain revealed that olfactory rods were present in the OE of *ift88*^−/−^ mutants at 5 dpf (Figure 3A, B).

**Figure 3.**
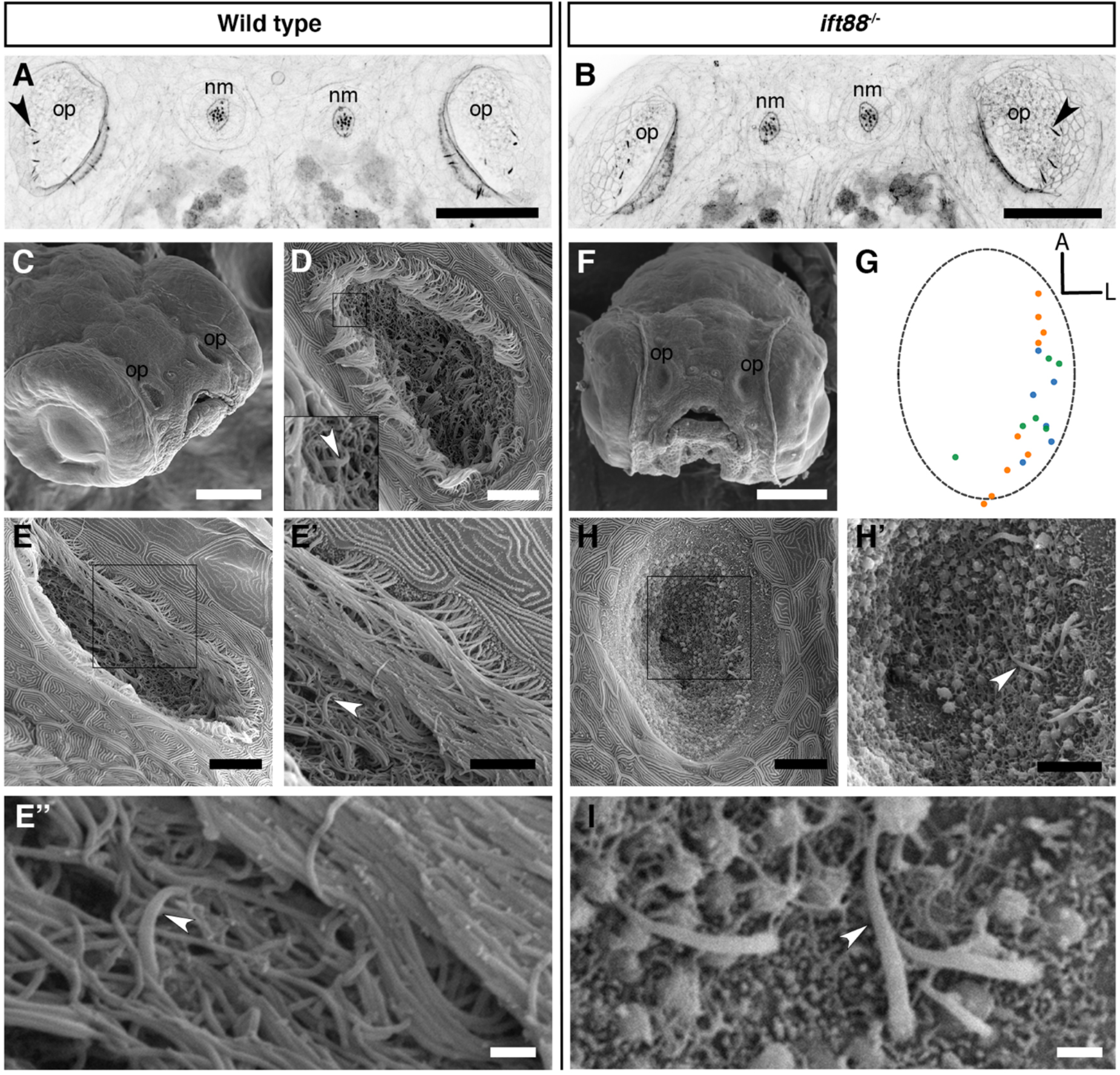
Olfactory rod cells are present in the olfactory epithelia of *ift88*^−/−^ zebrafish mutants, which lack cilia. (A, B) Maximum intensity projections of Airyscan confocal images of phalloidin stains of a 5 dpf wild-type (A) and *ift88*^−/−^ mutant (B) larva; dorsal views, anterior to the top. Grayscale values from the original fluorescence image have been inverted. Abbreviations: nm, cranial neuromast; op, olfactory pit. Several olfactory rods (arrowheads mark examples) are visible in each olfactory pit. Scale bar = 50 μm. (C) SEM of the head of a 4 dpf wild-type larva. Scale bar = 100 μm. (D-E) SEM of 4 dpf larval wild-type olfactory pits (enlarged from panel C). Scale bars = 10 μm. Insert in D shows enlarged view of boxed area in D. Arrowhead marks the tip of a rod cell apical projection surrounded by olfactory cilia. (E’) Enlarged view of boxed area in E. Arrowhead marks one olfactory rod. Scale bar = 5 μm. (E’’) Enlargement of olfactory rod in E’ (arrowhead). Scale bar = 1 μm. (F) Frontal view SEM of the head of a 4 dpf *ift88*^−/−^ mutant larva. Scale bar = 100 μm. (G) A map of the positions of olfactory rod cell projection emergence through the OE in *ift88*^−/−^ mutant larvae (*N* of olfactory pits = 3), based on SEM images at 4 dpf; anterior ‘A’ to the top, lateral ‘L’ to the right. One dot represents one olfactory rod. Different coloured dots represent rods from different larvae. (Compare with Figure 1E.) (H) SEM of 4 dpf larval *ift88*^−/−^ mutant olfactory pit (enlarged from panel F). Scale bar = 10 μm (H’) Enlarged view of boxed area in H. Arrowhead marks one example olfactory rod cell projection present despite the loss of cilia. Scale bar = 5 μm. (I) Enlarged SEM of olfactory rods (arrowhead marks example) in 4 dpf larval *ift88*^−/−^ mutant olfactory pit (from a different individual). Scale bar = 1 μm.

The absence of cilia in *ift88*^−/−^ mutants allowed us to examine morphology of the rods using scanning electron microscopy (SEM). In the phenotypically wild-type sibling OE, the rods were almost completely obscured by olfactory cilia, with only the occasional tip of a projection visible (Figure 3C-E’’). However, SEM images of the olfactory pit of *ift88*^−/−^ mutants at 4 dpf, which lack cilia, revealed the presence of rod-like projections with a similar size, number, smoothly tapering morphology, and spatial distribution to the actin-rich projections described above (Figure 3F-I). At their base, olfactory rods are wider in diameter (about 0.6 μm) than the olfactory cilia in wild-type larvae (0.2 μm in diameter, as is typical for many cilia). We conclude that olfactory rods can develop in the absence of cilia.

### Olfactory rods can be labelled in the live larva

To visualise olfactory rods in live larvae, we imaged the *Tg(actb2:Lifeact-RFP)* transgenic line at 4 and 6 dpf, and *Tg(actb2:Lifeact-GFP)* at 5 dpf (Behrndt et al., 2012). We found fluorescent apical projections in the olfactory pits of live larvae in all cases (*N* of fish = 4) (Figure 4A-C, Supplementary Movie 1). These matched the size, shape, and posterolateral distribution of rod cells present in fixed samples (Figure 4D, E). Despite potential shrinkage due to fixation, there was no overall difference in the lengths of projections between live and fixed samples (Figure 4E). The zig-zag pattern exhibited by RFP-positive olfactory rods in raster-scanned images of live larvae suggested that rods were moving during image capture (Figure 4B). Fast-capture time series imaging of the *Tg(actb2:Lifeact-RFP)* transgenic line allowed us to observe that the projection oscillates (Supplementary Movie 2), possibly as a result of ciliary beating.

**Figure 4.**
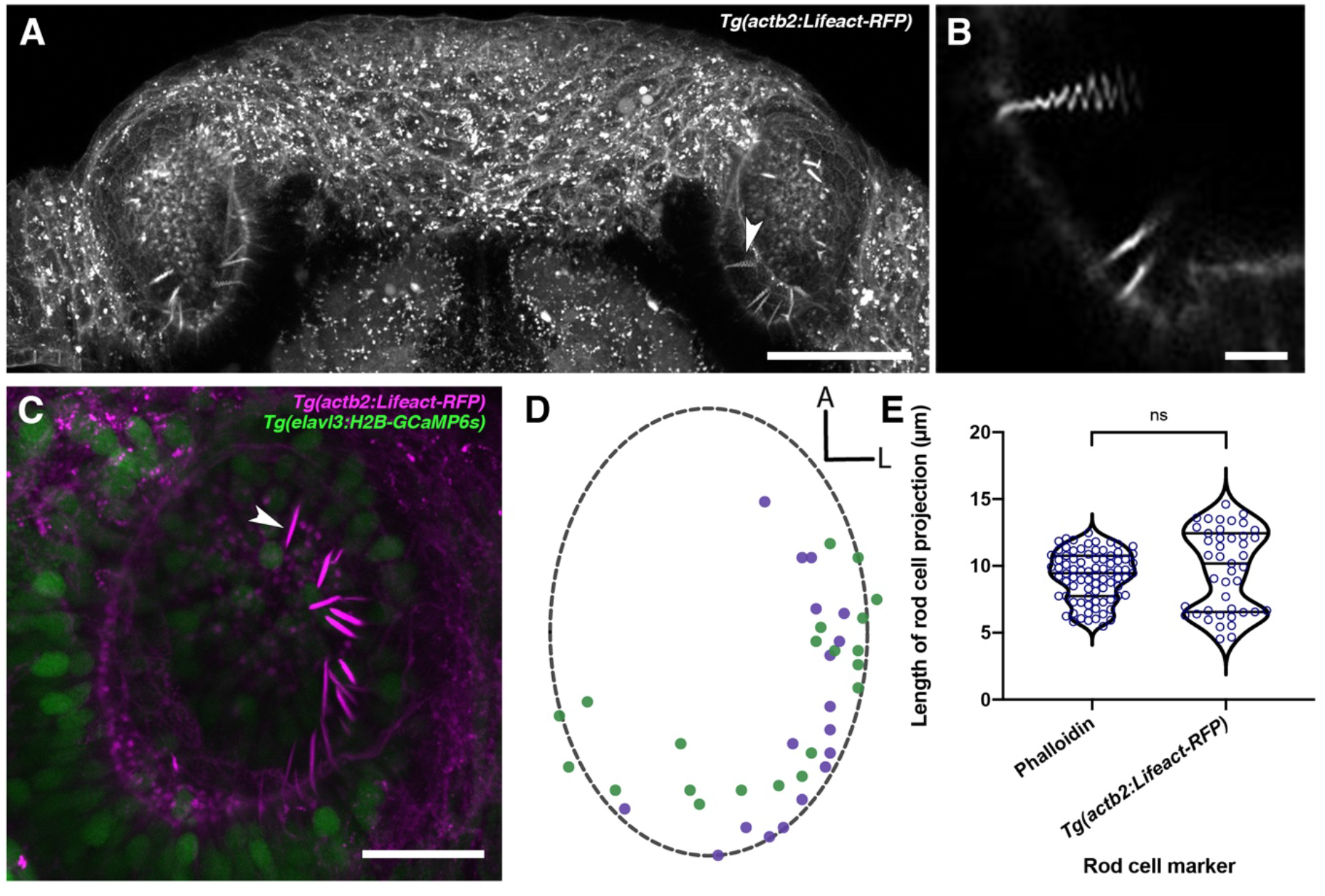
Olfactory rods are labelled in the olfactory epithelia of live zebrafish larvae by the *Tg(actb2:Lifeact-RFP)* transgene. (A) Maximum intensity projection of dorsal view image of the olfactory pits of a live 6 dpf *Tg(actb2:Lifeact-RFP)* transgenic larva; anterior to the top. Arrowhead marks one example olfactory rod positive for the Lifeact-RFP transgene. Scale bar = 50 μm. (B) Enlargement of olfactory rods in A (arrowhead in A) oscillating during raster-scanned image capture. (Raster scanning was performed from top to bottom in the image, as it has been rotated 90° clockwise.) (See Supplementary Movie 2.) Scale bar = 5 μm. (C) Maximum intensity projection image of a live 4 dpf *Tg(actb2:Lifeact-RFP);Tg(elavl3:H2B-GCaMP6s)* double-transgenic larval olfactory pit; anterior to the top, lateral to the right. Arrowhead marks one example olfactory rod positive for the Lifeact-RFP transgene (magenta). Neuronal nuclei are labelled in green. Larvae were fully mounted in agarose, so rods were not moving. Scale bar = 20 μm. (See Supplementary Movie 1.) (D) A map of the positions of olfactory rod cell projection bases in olfactory pits of 4 dpf *Tg(actb2:Lifeact-RFP);Tg(elavl3:H2B-GCaMP6s)* double-transgenic larvae (*N* of olfactory pits = 2), based on 2D maximum intensity projections of confocal images; anterior ‘A’ to the top, lateral ‘L’ to the right. One dot represents one olfactory rod. Different coloured dots represent rods from different larvae, with purple corresponding to panel C. (Compare with Figure 1E.) (E) A quantitative comparison of the lengths of olfactory rod cell projections in fixed larvae, using Alexa-phalloidin as a marker (*N* = 10, *n* of olfactory rods = 82) versus live larvae, using Lifeact-RFP as a marker (*N* = 2, *n* = 43). Violin plot; bars indicate the median and lower and upper quartiles for each group. Mann-Whitney U test; ns, not significant (*P* = 0.232).

### Neuronal promoters drive reporter expression in olfactory rod cells

To test whether rod cells are neuronal, we imaged two transgenic lines that have broad neuronal expression of cytoplasmic fluorescent reporters – *Tg(Xla.tubb:jGCaMP7f)* (Chia et al., 2019) (*N* of olfactory pits = 4) and *Tg(elavl3:GCaMP6f)* (Dunn et al., 2016) (*N* = 5). Dendrites and dendritic knobs of OSNs were clearly labelled by both lines. In some examples, we observed faintly-labelled projections extending from the surface of the olfactory epithelium, with a similar length and morphology to olfactory rods (Figure 5A-B). Imaging of double-transgenic *Tg(elavl3:GCaMP6f);Tg(actb2:Lifeact-RFP)* larvae at 5 dpf indicated that rods were GCaMP6f-positive (*N* of fish = 3; Figure 5C-C’’). These observations suggest that olfactory rod cells may be neurons.

**Figure 5.**
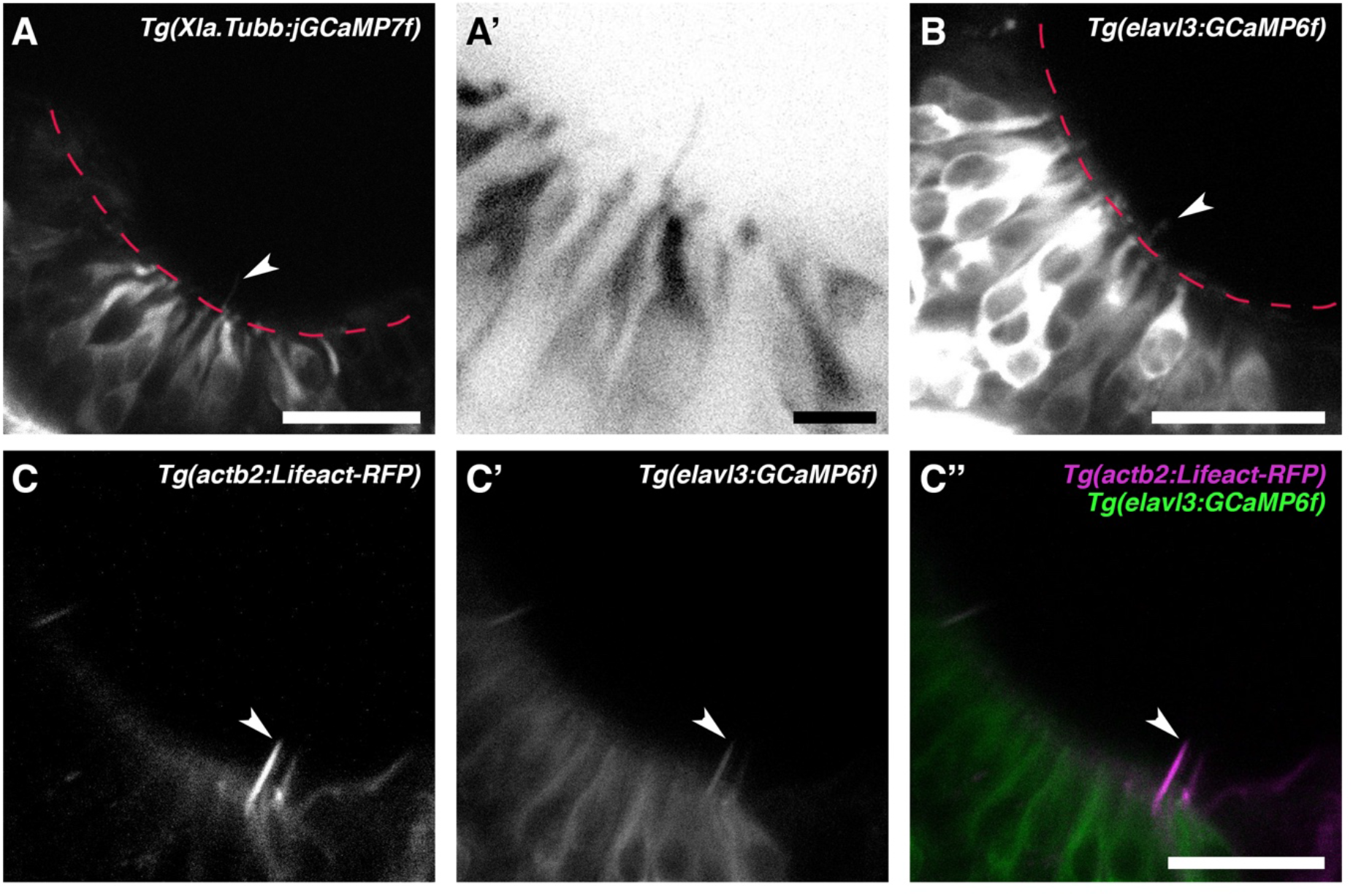
Olfactory rod cells are labelled by the cytoplasmic neuronal markers *Tg(Xla.Tubb:jGCaMP7f)* and *Tg(elavl3:GCaMP6f)*. (A) Olfactory pit of a 4 dpf *Tg(Xla.Tubb:jGCaMP7f)* larva; anterior to the top, lateral to the right. Red dotted line outlines the apical surface of the OE; arrowhead marks one olfactory rod, albeit faintly labelled. Scale bar = 20 μm. (A’) Enlargement of olfactory rod marked by arrowhead in A (grayscale values inverted). Scale bar = 5 μm. (B) Olfactory pit of a 5 dpf *Tg(elavl3:GCaMP6f)* larva; anterior to the top, lateral to the right. Red dotted line outlines the apical surface of the OE; arrowhead marks one example olfactory rod, albeit faintly labelled. Scale bar = 20 μm. (C-C’’) Lifeact-RFP signal (C), GCaMP6f signal (C’), and merged signals (C’’) in an olfactory pit of a 5 dpf *Tg(elavl3:GCaMP6f);Tg(actb2:Lifeact-RFP)* double-transgenic larva; anterior to the top, lateral to the right. Arrowhead marks one example olfactory rod, positive for both Lifeact-RFP and GCaMP6f. Scale bar = 20 μm.

### Olfactory rod cells are not hair-cell-like cells

Given the superficial similarity in appearance of the olfactory rod to hair-cell stereocilia in phalloidin stains, and a report of a rare cell type bearing stereocilia-like microvilli in the rat OE (Menco & Jackson, 1997), we tested whether there is any similarity between olfactory rod cells and mechanosensory hair cells of the inner ear and lateral line. As shown in Figures 1 and 3, the zebrafish olfactory rod appears to be a single structure rather than a collection of microvilli or stereocilia. To test whether olfactory rod cells express sensory hair cell markers, we performed an Alexa-phalloidin co-stain on the *Tg(pou4f3:GFP)* transgenic line, a known marker for hair cells (Xiao et al., 2005). At 5 dpf, the stereociliary bundle of lateral line neuromast hair cells was clearly marked by both GFP and phalloidin, which acted as our positive control (Figure 6A-A’’). However, the GFP did not co-localise with the phalloidin signal in the olfactory rods, or in the cell body beneath a phalloidin-positive rod (Figure 6B-B’’). Additionally, confocal images of an antibody stain against synaptic vesicle protein 2 (SV2), a marker of synaptic terminals of afferent neurons contacting secondary sensory cells such as mechanosensory hair cells and taste receptors (Buckley & Kelly, 1985; Portela-Gomes et al., 2000; Zachar & Jonz, 2012), showed that SV2-positive synapses are not present in the olfactory epithelia at 4 dpf (Figure 6C, D). This contrasted with strong staining of synaptic terminals in cranial lateral line neuromasts in the vicinity of the olfactory pits (Fig. 6C, D).

**Figure 6.**
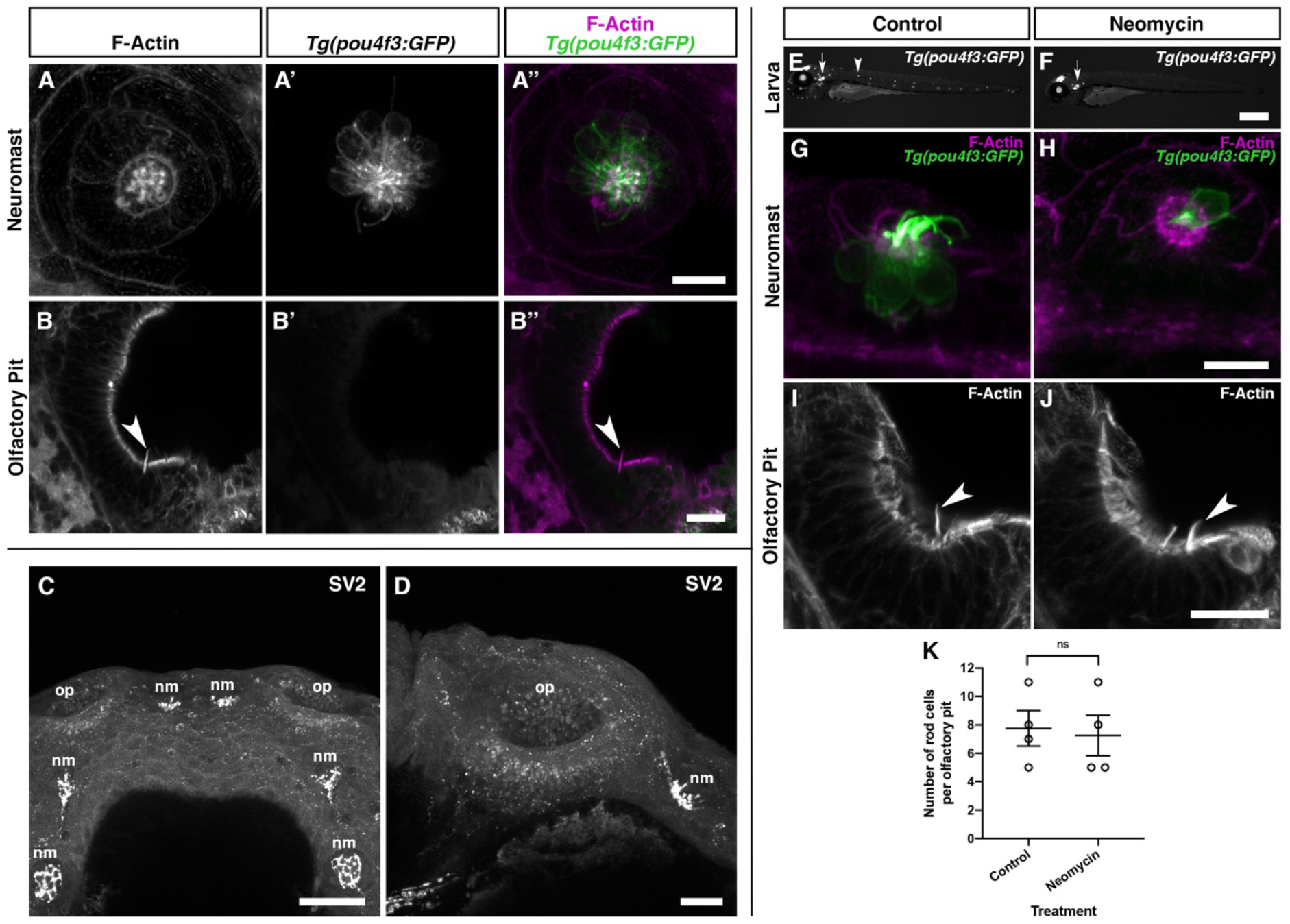
Olfactory rod cells in the zebrafish olfactory epithelium are not hair-cell-like. (A-A’’) Maximum intensity projection of Airyscan confocal image of Alexa-phalloidin signal (A), *Tg(pou4f3:GFP)* signal (A’), and merged signals (A’’) in a cranial neuromast of a 5 dpf larva. Scale bar = 10 μm. (B-B’’) Airyscan confocal image of Alexa-phalloidin signal (B), *Tg(pou4f3:GFP)* signal (B’), and merged signals (B’’) in an olfactory pit of a 5 dpf larva; anterior to the top, lateral to the right. Arrowhead marks one olfactory rod. Scale bar = 20 μm. (C-D) Maximum intensity projections of confocal images of synaptic vesicle 2 (SV2) protein immunohistochemistry signal in the heads of 4 dpf wild-type larvae. Abbreviations: nm, cranial neuromast; op, olfactory pit. (C) Anterior to the top. Scale bar = 50 μm. (D) Anterior to the top, lateral to the right. Scale bar = 20 μm. (E-F) Widefield imaging of 3 dpf *Tg(pou4f3:GFP)* larvae showing the damaging effects of 500 μM neomycin treatment for 60 minutes on lateral line neuromast hair cells. Fluorescence is lost or greatly reduced in both trunk (arrowhead) and cranial neuromasts, whereas fluorescence in hair cells of the inner ear maculae and cristae (arrow) is unaffected. Scale bar = 500 μm. (G-H) Maximum intensity projections of Airyscan confocal images showing the damaging effects of 500 μM neomycin treatment for 60 minutes on hair cells in a cranial neuromast of a 3 dpf larva, using *Tg(pou4f3:GFP)* (green) and Alexa-phalloidin (magenta) as markers. Scale bar = 10 μm. (I-J) Maximum intensity projections of Airyscan confocal images showing no effect of 500 μM neomycin treatment for 60 minutes on olfactory rods, using Alexa-phalloidin as a marker; anterior to the top, lateral to the right. Arrowheads mark olfactory rods. Scale bar = 20 μm. (K) The number of olfactory rod cell projections per olfactory pit of 3 dpf *Tg(pou4f3:GFP)* larvae after 500 μM neomycin treatment for 60 minutes (*N* of olfactory pits = 4), compared with an untreated group (*N* = 4). Welch’s unpaired two-tailed *t*-test; ns, not significant (*P* = 0.8018).

We next investigated whether treatment with neomycin, an aminoglycoside antibiotic and well-described ototoxin, has the same damaging effect on olfactory rod cells as on lateral line hair cells (Harris et al., 2003). Following neomycin treatment at 500 μM for 60 minutes on 3 dpf *Tg(pou4f3:GFP)* larvae, lateral line hair cells were lost or severely damaged, as determined by a decrease in the number of GFP-positive cells in both cranial and trunk neuromasts and a change in morphology of any remaining cells (Figure 6E-H). By contrast, olfactory rods appeared unaffected (Figure 6I, J), with no significant change in the number of rods present in each olfactory pit (Figure 6K). Taken together, the smooth appearance of the olfactory rods, lack of hair cell and synaptic vesicle marker expression, and resistance to neomycin indicate that olfactory rod cells are not closely related to hair cells.

### A sub-population of olfactory rod cells expresses a Lifeact transgene driven by the sox10 promoter

Sox10 is a known marker of both neural crest and otic epithelium (Dutton et al., 2001). Robust transgene expression driven by the *sox10* promoter has been reported in the OE and other tissues in the zebrafish (Mongera et al., 2013; Saxena et al., 2013). We have generated a *Tg(sox10:Lifeact-mRFPruby)* transgenic line to visualise actin localisation and dynamics in the live embryo in *sox10*-expressing tissues. As reported for the *Tg(sox10:eGFP)* transgene (Saxena et al., 2013), we observed OSNs expressing *Tg(sox10:Lifeact-mRFPruby)* in the OE at 4 and 5 dpf; based on morphology, most of these cells were microvillous neurons. However, staining with Alexa-phalloidin on fixed samples revealed the co-expression of Lifeact-mRFPruby in a sub-population of phalloidin-positive olfactory rod cell projections (Figure 7A-B’’). Not all olfactory rod cells expressed the transgene; an average of 64.4% of rod cells marked by phalloidin (*N* of olfactory pits = 5, *n* of olfactory rods = 59) also expressed Lifeact-mRFPruby (Figure 7C). As for the rods labelled with Lifeact-RFP, rods labelled with Lifeact-mRFPruby oscillated (Supplementary Movie 3).

**Figure 7.**
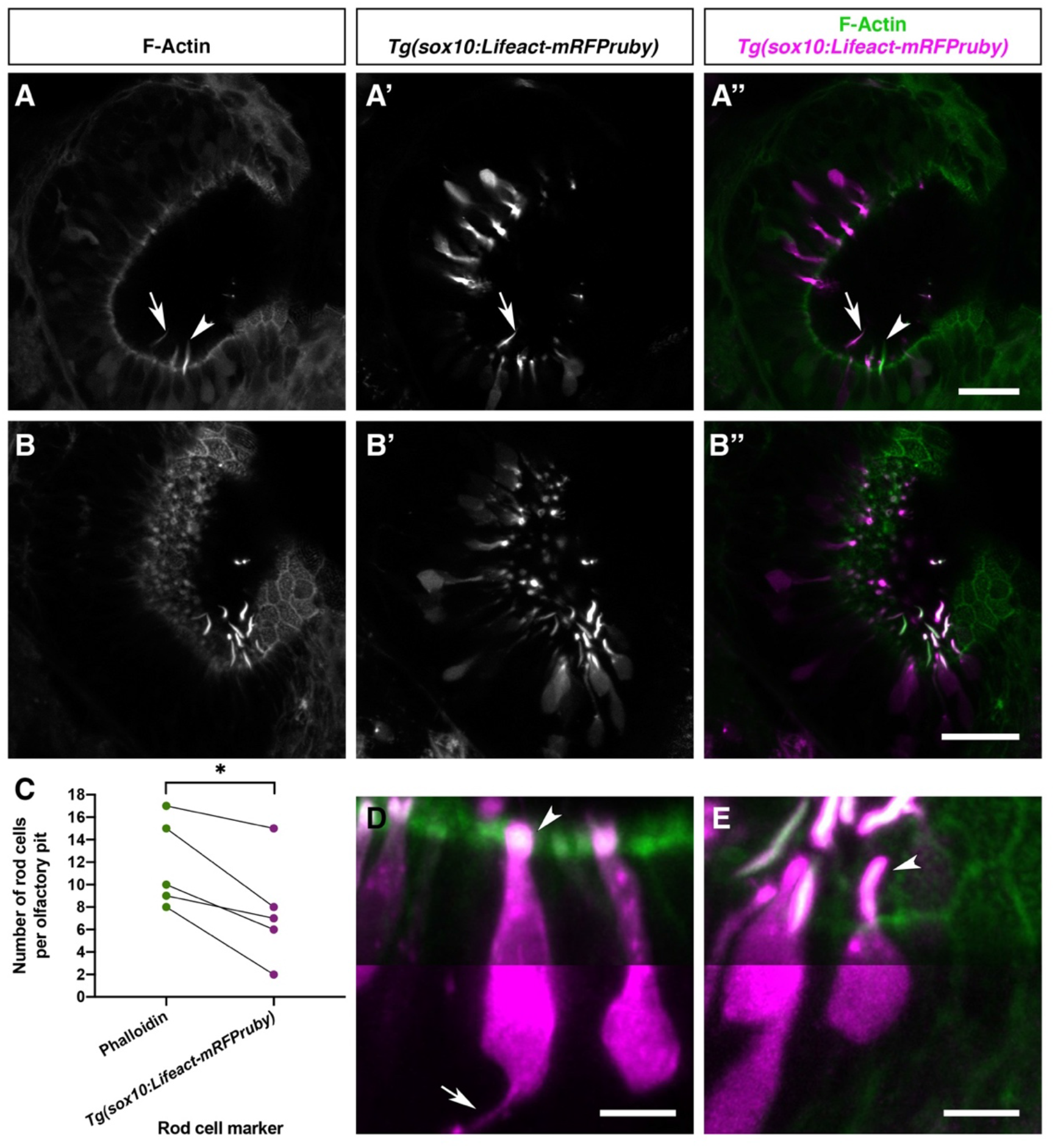
Olfactory rod cells are apically located in the zebrafish olfactory epithelium, with a rounded cell body and no detectable axon. (A-B’’) Airyscan confocal image of Alexa-phalloidin signal (A, B), *Tg(sox10:Lifeact-mRFPruby)* signal (A’, B’), and merged signals (A’’, B’’) in olfactory pits of 4-5 dpf larvae; anterior to the top, lateral to the right. Arrowhead marks one olfactory rod negative for Lifeact-mRFPruby. Arrow marks one olfactory rod positive for Lifeact-mRFPruby. Scale bars = 20 μm. (C) Number of olfactory rod cells positively marked by Alexa-phalloidin (*n* of olfactory rods = 59), compared with the number of those also marked by *Tg(sox10:Lifeact-mRFPruby)* (*n* = 38), in olfactory pits of 4-5 dpf larvae (*N* of olfactory pits = 5). Connecting lines indicate rods from the same olfactory pit. Paired two-tailed *t*-test; * indicates *P* = 0.0146. (D) Enlargement of two microvillous OSNs, expressing Lifeact-mRFPruby, in the OE of a 4 dpf larva; Alexa-phalloidin signal (green), *Tg(sox10:Lifeact-mRFPruby)* signal (magenta). Arrowhead marks the microvillous apical projections. The gamma value for the magenta channel in the bottom half of the panel has been adjusted to show the axon from one of the cells (arrow). Scale bar = 5 μm. (E) Enlargement of olfactory rod cells (of which both the apical actin projections and cell bodies are labelled by the *Tg(sox10:Lifeact-mRFPruby)* transgene) in the OE of a 4 dpf larva; Alexa-phalloidin signal (green), *Tg(sox10:Lifeact-mRFPruby)* signal (magenta). Arrowhead marks a rod cell apical projection, positive for both markers. The gamma value for the bottom half of the panel has been adjusted as in D; no axon is visible. Scale bar = 5 μm. See also Supplementary Movie 3.

The sparse expression of the *Tg(sox10:Lifeact-mRFPruby)* transgene allowed us to visualise the morphology of the cell body of olfactory rod cells and ask whether they have an axon. Lifeact-mRFPruby-expressing cell bodies were positioned apically in the OE and were rounded in shape (Figure 7B-B’’, E). They were morphologically distinct from the well-described microvillous neurons (Figure 7D, E) as well as ciliated and crypt OSNs. The axons of microvillous OSNs were visible in those cells labelled by the transgene (Figure 7D). However, with this marker, we were unable to observe an axon extending from the cell body of olfactory rod cells (*N* of olfactory pits = 5, *n* of cells = 9; Figure 7E).

To test whether the development of olfactory rod cells is dependent on *sox10* function, we stained *sox10*^−/−^ homozygous mutants (Dutton et al., 2001) with Alexa-phalloidin. Olfactory rods were present in *sox10*^−/−^ mutants at 5 dpf, but variable in number (*N* of olfactory pits = 8, *n* of olfactory rods = 53; Figure 8). Taken together, the data from *Tg(sox10:Lifeact-mRFPruby)* transgenic and *sox10*^−/−^ mutant larvae indicate that *sox10* function is not essential for the formation of olfactory rods.

**Figure 8.**
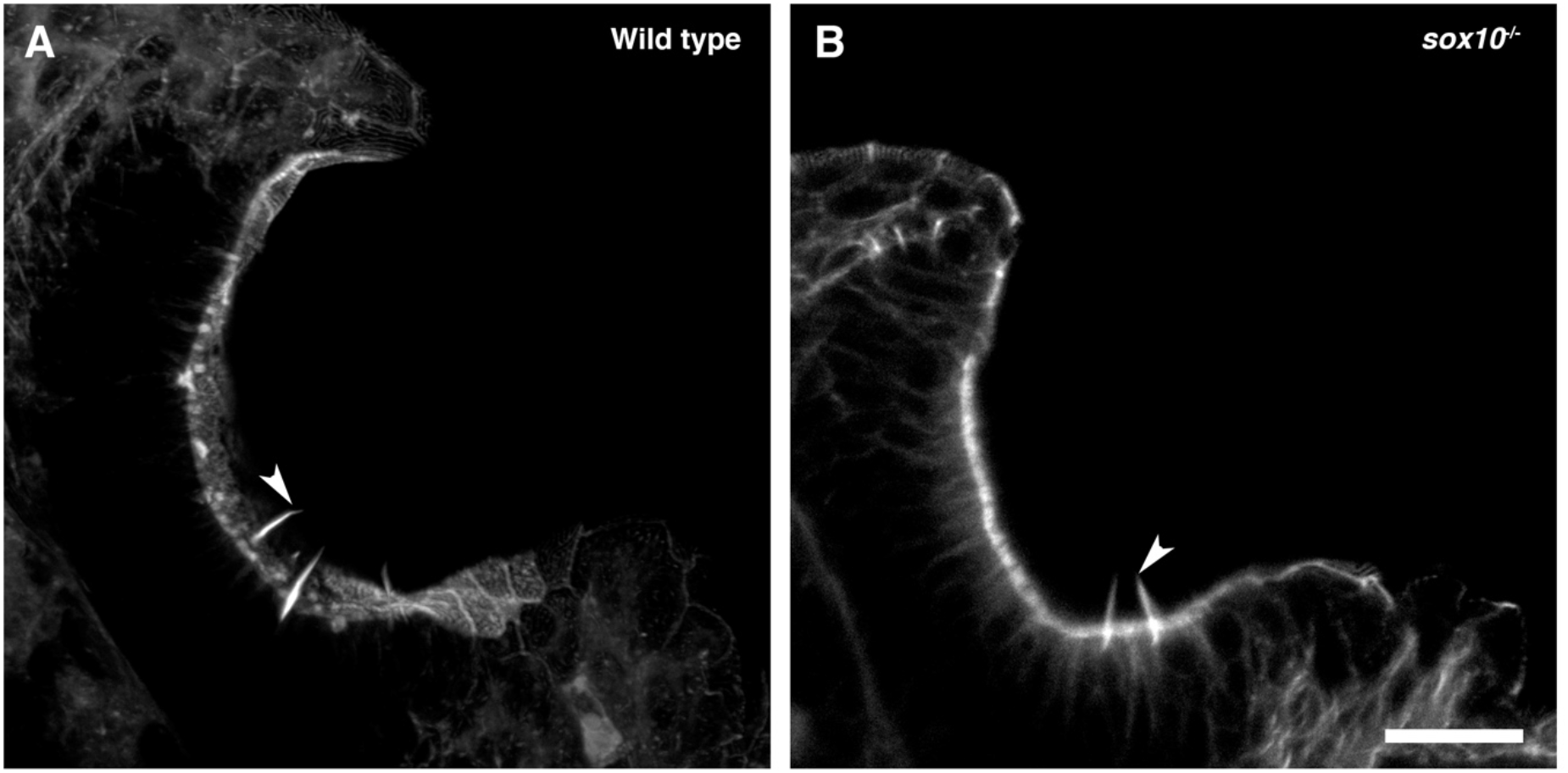
Olfactory rod cells are present in the olfactory epithelia of *sox10*^−/−^ zebrafish mutants. (A) Maximum intensity projection of Airyscan confocal image of phalloidin stain in a 5 dpf larval wild-type olfactory pit; anterior to the top, lateral to the right. Arrowhead marks one example olfactory rod. Scale bar = 20 μm. (B) Airyscan confocal image of phalloidin stain in a 5 dpf larval *sox10*^−/−^ mutant olfactory pit; anterior to the top, lateral to the right. Arrowhead marks one example olfactory rod. Scale bar = 20 μm.

## Discussion

Through the use of phalloidin staining, immunohistochemistry, transgenic zebrafish lines, SEM and high-resolution fluorescence confocal imaging, we have identified a rare cell type, the olfactory rod cell, in the zebrafish larval and juvenile OE. Olfactory rod cells, which have not previously been described in zebrafish to our knowledge, are morphologically distinct from the well-characterised OSNs and other known cell types in terms of their apical projections, cell shape, and distribution and positioning within the OE.

### The olfactory rod: an actin-rich apical projection

The spectacular actin-rich projection of the olfactory rod cell adds to the rich repertoire of known F-actin-based cellular specialisations, which include microvilli, stereocilia, lamellipodia, filopodia, cytonemes and microridges (reviewed in [Heath & Holifield, 1991; Theriot & Mitchison, 1991]; Ramírez-Weber & Kornberg, 1999; Pinto et al., 2019; Inaba et al., 2020). Many classes of sensory cell, in both fish and mammals, bear actin-rich mechano-or chemosensory microvillous projections, including the stereocilia of sensory hair cells (Tilney et al., 1980; reviewed in [Gillespie & Müller, 2009; Barr-Gillespie, 2015]), and the microvilli of olfactory and vomeronasal microvillous neurons, solitary chemosensory cells (SCCs) of the skin and barbel (Kotrschal et al., 1997; Finger et al., 2003), taste bud cells (Hansen et al., 2002; Zachar & Jonz, 2012), spinal cerebrospinal fluid-contacting neurons (CSF-cNs; Djenoune et al., 2014; Desban et al., 2019), Merkel cells, retinal Müller glia (Sekerková et al., 2004), and the brush and tuft cells of mammalian respiratory and intestinal epithelia, respectively (reviewed in [Reid et al., 2005; Schneider et al., 2019]). As a single structure with a smoothly tapering morphology, the zebrafish olfactory rod differs from these oligovillous structures. Adult zebrafish SCCs, found distributed over the entire body surface (Kotrschal et al., 1997), and mature light cells of the zebrafish taste bud (Hansen et al., 2002) each bear a single microvillus, but at 1-3 μm in length, these are much shorter than the olfactory rods we describe.

A note on terminology: olfactory rod cells are distinct from rodlet cells, which have been reported in many different epithelial tissues (including OE) of marine and freshwater fish, including zebrafish, and contain several intracellular electron-dense rodlets within a thick cuticular-like wall (Bannister, 1966; reviewed in [Morrison & Odense, 1978]; Hansen & Zeiske, 1998 Dezfuli et al., 2007; DePasquale, 2020). Recently, phalloidin staining has demonstrated that the rodlets, which can be extruded from the cell, are not composed of F-actin (DePasquale, 2020). Thus, zebrafish olfactory rod cells, which are unique to the OE at the larval stages we have described, are not related to rodlet cells.

### Olfactory rod cells in other teleost species

Previous studies have provided brief descriptions of similar cell types in other teleost species, including the common minnow (Bannister, 1965), several eel species (Schulte, 1972; Yamamoto & Ueda, 1978), goldfish (Breipohl et al., 1973; Ichikawa & Ueda, 1977), rainbow trout (Rhein et al., 1981), common bleak (Hernádi, 1993), catfish (Datta & Bandopadhyay, 1997), and several cave fish and cave loach species (Waryani et al., 2013; Waryani et al., 2015; Zhang et al., 2018).

Using transmission electron microscopy (TEM), Bannister (1965) reported sparsely-populated rod-shaped protrusions, approximately 4 μm in length and shorter than surrounding sensory and non-sensory olfactory cilia, in the OE of adult (3.7 cm) common minnow (*Phoxinus phoxinus*). Here, the rod-like projection consisted of several bundles of fibres, consistent with the appearance of F-actin, extending from deep within the cell (Bannister, 1965). Similarly, using TEM and SEM respectively, Schulte (1972) and Yamamoto and Ueda (1978) reported the presence of olfactory rod cells in the OE of several adult eel species: European eel (*Anguilla anguilla*), Japanese eel (*A. japonica*), white-spotted conger (*Conger myriaster*), buffoon snake eel (*Microdonophis erabo*), and brutal moray (*Gymnothorax kikado*). In European eels, the cells were described as a receptor with a single rod-shaped appendage, measuring 0.8 μm in diameter and extending 4 μm above the apical surface of the epithelium (Schulte, 1972). Olfactory rods in the other four species measured 1 μm in diameter and 10 μm in length. Rods were either found to exist solitarily or in a group; interestingly, it was noted that olfactory cilia were sparse in areas where rods occurred in a group (Yamamoto & Ueda, 1978).

More recent reports include comparisons of the surface structures of olfactory epithelia in different adult cave fish and loaches. SEMs in *Sinocyclocheilus jii* and *S. furcodorsalis* cave fish, and in *Oreonectes polystigmus* and *O. guananensis* cave loaches revealed that olfactory rods were clustered in different regions of olfactory rosette lamellae (Waryani et al., 2013; Waryani et al., 2015). Another SEM study on the variations in olfactory systems of adult cave fish species of different habitats reported not just one, but three different cell types all classified as ‘rod cilia’ in the olfactory epithelia of *S. anshuiensis* and *S. tianlinensis*. The first cell type had a long base with an oval apex, the second contained an oval base with a thin apex, while the third was rod-shaped and thin from base to tip, measuring 2.01-3.08 μm in length (Zhang et al., 2018). Despite the shorter length, this third type appeared morphologically consistent with zebrafish olfactory rod cells. Unlike other teleosts, olfactory rod cells were reported as the dominant cell type over ciliated and microvillous neurons in the OE of *S. jii* (Waryani et al., 2013). This may be an example of the known compensatory enhancement of the olfactory system in blind morphs of cave fish (Bibliowicz et al., 2013; reviewed in [Krishnan & Rohner, 2017]).

Although there appear to be variations in the numbers and sizes of olfactory rod cells reported in these other teleost species, some of these cells may be homologous to the olfactory rod cells we describe in zebrafish larvae. However, all of these previous studies were limited to fixed adult samples by means of TEM and SEM, and none have tested or confirmed the molecular composition of the rod.

### Olfactory rod cells differ from known olfactory sensory neurons or secondary receptor cells

We have detected weak expression of cytoplasmic fluorescent markers driven by neuronal promoters in olfactory rod cells, suggesting they are a type of OSN. However, we were unable to detect an axon in nine individual olfactory rod cells imaged with a Lifeact-mRFPruby transgene at 4-5 dpf. Of note, Ichikawa and Ueda (1977) performed olfactory nerve bundle transection in adult goldfish to determine which cell types are olfactory receptors. As expected, transection caused retrograde degeneration of both ciliated and microvillous OSNs. Olfactory rod cells, however, were still identifiable by SEM in the OE 10 days after nerve transection. The authors concluded that adult goldfish olfactory rod cells are not sensory receptor cells; however, the result could also indicate differences in regeneration kinetics between different neuronal cell types.

The shape, position and actin-rich projection of zebrafish olfactory rod cells, however, do have some similarities to those of kappe OSNs, identified to date only in adult zebrafish OE (Ahuja et al., 2014). The actin-positive apical specialisation of kappe cells differs in morphology from the rods we describe, and is thought to consist of microvilli, although this has not been confirmed at an ultrastructural level. Kappe neurons were reported to have an axon, and were thus interpreted as a class of OSN (Ahuja et al., 2014). It will be important to determine whether olfactory rod cells are present in the adult zebrafish OE and how their morphology relates to kappe cells.

Olfactory rod cells do not appear to share characteristics with sensory hair cells. As a monovillous structure, the olfactory rod is quite unlike the stereociliary bundle of a hair cell; moreover, the cells do not express known hair cell markers, and are insensitive to the aminoglycoside antibiotic neomycin, a potent ototoxin. In addition, the entire olfactory pit was negative for the synaptic vesicle and neuroendocrine cell marker SV2 (Buckley & Kelly, 1985; Portela-Gomes et al., 2000), indicating that olfactory rod cells are unlikely to be secondary sensory cells innervated by afferent neurons.

### Olfactory rod cells as artefact

Since the first report of olfactory rod cells, several studies have proposed that they may represent senescent forms of OSNs or fixation artefacts (Muller & Marc, 1984; Moran et al., 1992; reviewed in [Hansen & Zielinski, 2005]). A study in the goldfish (*Carassius auratus*) and channel catfish (*Ictalurus punctatus*), using TEM, SEM and filling with horseradish peroxidase, concluded that rods are most likely a result of fusion of olfactory cilia or microvilli – an indicator of ageing OSNs (Muller & Marc, 1984). A later study on the ultrastructure of olfactory mucosa in brown trout (*Salmo trutta*) also classified rods as products of the fusion of olfactory cilia during fixation (Moran et al., 1992). Indeed, TEM images in this study showed multiple ciliary axonemes surrounded by a single membrane (Moran et al., 1992). The presence of such fixation artefacts has led to frequent dismissal of olfactory rod cells in the literature, for example in juvenile and adult European eels (Sola et al., 1993). In the zebrafish, however, the olfactory rods we describe are clearly not a fixation artefact, as they are present in the live larva. Moreover, they are not formed by fusion of cilia, as the olfactory rods are F-actin-positive, do not stain with an anti-acetylated α-tubulin antibody, and are present in *ift88*^−/−^ mutants, which lack cilia.

### Possible functions of olfactory rod cells

Actin-rich projections on sensory cells are known to have mechanosensory (reviewed in [Gillespie & Müller, 2009]), chemosensory (Höfer & Drenckhahn, 1999; Hansen et al., 2002; Zachar & Jonz, 2012), or multimodal functions (for example in CSF-cNs in zebrafish; Djenoune et al., 2014; Desban et al., 2019). A mechanosensory role for zebrafish olfactory rod cells, for example in detecting ciliary movement or ciliary-driven fluid flow, or a chemosensory role in detecting odorants, could aid olfactory perception in the larva. The olfactory rod cell has previously been referred to as a sensory receptor cell in studies of other teleost species (Bannister, 1965; Schulte, 1972; Breipohl et al., 1973; Waryani et al., 2013; Waryani et al., 2015). Although functions were not explicitly investigated, Bannister (1965) speculated that based on their internal structure, minnow olfactory rod cells perform specific chemosensory roles. Another possibility is that they could correspond to brush or tuft cells in air-breathing mammals, which have important roles in immunity (Andres, 1975; reviewed in [Reid et al., 2005]; Howitt et al., 2016; reviewed in [Schneider et al., 2019]). These ideas remain to be tested.

### Possible origins of olfactory rod cells

Our work does not address the developmental origin of olfactory rod cells, but it is of interest that they express a *sox10*-driven transgene, albeit in a mosaic fashion. *Sox10* mRNA is frequently described as a neural crest marker, but is also expressed strongly in otic epithelium (Dutton et al., 2001), a placodally-derived tissue. The use of *sox10*-driven transgenic lines to identify neural crest derivatives remains controversial. Expression of a *sox10*:eGFP transgene together with photoconversion studies has led to the conclusion that a subpopulation of microvillous neurons in the OE is derived from neural crest (Saxena et al., 2013), and use of an inducible *sox10*:*ER*^*T2*^-Cre transgenic line has identified previously ‘contested’ neural crest derivatives, including cells in the sensory barbels (Mongera et al., 2013). However, using lineage reconstruction through backtracking and photoconversion experiments, Aguillon et al. (2018) have argued that all olfactory neurons, including OSNs and gonadotropin-releasing hormone 3 (GnRH3) cells, are derived entirely from preplacodal progenitors. We think it likely that olfactory rod cells are placodally-derived, but this remains to be confirmed.

The *Tg(sox10:Lifeact-mFRPruby)* line is expressed in a subset of both olfactory rod cells and of microvillous neurons, with variation in the proportion of expressing cells between individuals. This could reflect true heterogeneity in the olfactory rod cell and microvillous neuron populations, or it could be a result of mosaic or leaky expression of the transgene. Mosaic expression is typical for many transgenes (Mosimann et al., 2013), while leaky expression, which can be explained through the lack of appropriate silencer elements (Jessen et al., 1999), is suspected for the *sox10* promoter fragment used in our transgenic construct (reviewed in [Tang & Bronner, 2020]). Nevertheless, the *Tg(sox10:Lifeact-mRFPruby)* line has proved a fortuitous tool for visualising olfactory rod cells in the live larva.

### Concluding remarks

As a key model organism for the study of the olfactory system (reviewed in [Kermen et al., 2013; Calvo-Ochoa & Byrd-Jacobs, 2019]), a complete inventory of the cell types present in the zebrafish OE will be an important resource and reference point for further study. Olfactory dysfunction can signify underlying cellular disorders and can also be implicated in neurodegenerative diseases (reviewed in [Whitlock, 2015]; Bergboer et al., 2018). OSNs project directly to the OB, and thus provide an entry route for pathogens to the brain (reviewed in [Dando et al., 2014]). Cells in the OE can themselves be damaged by viral infection, leading to a reduction, change, or loss of sense of smell, a phenomenon that has attracted much recent attention due to the damaging action of SARS-CoV-2 on the human olfactory system (Brann et al., 2020; Gupta et al., 2020). In the zebrafish, new functions of the OE, such as the detection of sodium and chloride ions (Herrera et al., 2020), continue to be uncovered. Addressing the functions of the olfactory rod cell will be an important next step.

## Ethics

All zebrafish work in Sheffield was undertaken under licence from the UK Home Office and according to recommended standard husbandry conditions (Aleström et al., 2019). All experiments in Singapore were performed under guidelines approved by the Institutional Animal Care and Use Committee of Biopolis.

## Author Contributions

Designed the research: KYC, TTW, SJJ. Conducted the experiments: KYC, SJJ, TTW, SB, NJvH, MM, CJH. Data analysis: KYC, SJJ, TTW. Writing (original draft): KYC, TTW; writing (review and editing): KYC, TTW, SJJ, with additional contributions from SB, NJvH, CJH.

## Acknowledgements

We thank Karen Carmargo Sosa and Robert Kelsh for providing fixed *sox10*^−/−^ larvae. We thank Henry Roehl for making the p5E *-4725 sox10* promoter (originally from the Kelsh lab), *Lifeact-mRFPruby* construct (originally from the Wedlich-Söldner and Sixt labs), and Zeiss Axio Zoom.V16 microscope available to us, Ana Almeida Jones for help with imaging, Emily Glendenning for technical support, and members of the Whitfield lab for discussion. We are grateful to the Sheffield Aquarium Team for excellent fish care. We also thank Kathleen Cheow, Ruey-Kuang Cheng, Jason Lai, and Tim Saunders for assistance with fish in Singapore.

## Supplementary Material

**Supplementary Movie 1. Olfactory rods are labelled in the olfactory epithelia of live zebrafish by the *Tg(actb2:Lifeact-RFP)* transgene.**

3D rendering of a confocal image of a 4 dpf *Tg(actb2:Lifeact-RFP)*;*Tg(elavl3:H2B-GCaMPs)* double-transgenic larval olfactory pit; anterior to the top. Olfactory rods are labelled in magenta; neuronal nuclei are labelled in green.

**Supplementary Movie 2. Olfactory rods labelled with Lifeact-RFP in the olfactory epithelia of live zebrafish larvae oscillate.**

Fast-capture time series confocal imaging (4.35 fps) of olfactory rods in a 6 dpf *Tg(actb2:Lifeact-RFP)* larva; anterior to the top, lateral to the right. Playback speed of the movie is 4 fps. Scale bar = 20 μm.

**Supplementary Movie 3. Olfactory rods labelled with Lifeact-mRFPruby in the olfactory epithelia of live zebrafish larvae oscillate.**

Fast-capture time series light-sheet imaging (50.04 fps) of a 5 dpf *Tg(sox10:Lifeact-mRFPruby)* larval olfactory pit; anterior to the top left, lateral to the top right. Beating olfactory cilia are visible in brightfield (grayscale), and oscillating olfactory rods are labelled by Lifeact-mRFPruby (magenta). Playback speed of the movie is 7 fps. Scale bar = 20 μm.

